# Metabolome and transcriptome analysis reveals components regulating triterpenoid saponin biosynthesis of soapberry

**DOI:** 10.1101/2022.02.28.482332

**Authors:** Yuanyuan Xu, Guochun Zhao, Xiangqin Ji, Jiming Liu, Tianyun Zhao, Yuan Gao, Shilun Gao, Yingying Hao, Yuhan Gao, Lixian Wang, Xuehuang Weng, Zhong Chen, Liming Jia

## Abstract

Soapberry (*Sapindus mukorossi* Gaertn.) pericarps are rich in valuable bioactive triterpenoid saponins. However, the saponin content dynamics and the molecular regulatory network of saponin biosynthesis in soapberry pericarps remain largely unclear. Here, we performed combined metabolite profiling and transcriptome analysis to identify saponin accumulation kinetic patterns, investigate gene networks, and characterize key candidate genes and transcription factors involved in saponin biosynthesis in soapberry pericarps. A total of 54 saponins were tentatively identified, including 25 that were differentially accumulated. Furthermore, 49 genes putatively involved in sapogenin backbone biosynthesis and some candidate genes assumed to be responsible for the backbone modification, including 41 cytochrome P450s and 45 glycosyltransferases, were identified. Saponin-specific clusters/modules were identified by Mfuzz clustering and weighted gene co-expression network analysis, and one TF–gene regulatory network underlying saponin biosynthesis was proposed. The results of yeast one-hybrid assay and electrophoretic mobility shift assay suggested that SmbHLH2, SmTCP4, and SmWRKY27 may play important roles in the triterpenoid saponin biosynthesis by directly regulating the transcription of *SmCYP71D-3* in soapberry pericarp. Overall, these findings provide valuable information for understanding the molecular regulatory mechanism of saponin biosynthesis, enriching the gene resources, and guiding further research on triterpenoid saponin accumulation in soapberry pericarps.

**One–sentence summary:** Combining metabolome and transcriptome analysis to identify saponin kinetic patterns, gene networks, and key candidate genes and transcription factors involved in saponin biosynthesis of soapberry.

## Introduction

Soapberry (*Sapindus mukorossi* Gaertn.) is an economically important tree belonging to the Sapindaceae that grows in southern China, the Indochina Peninsula, India, Japan and North Korea (Jia and Sun, 2012; Sun et al., 2017). The tree has been used in traditional medicine for centuries in China because its pericarps possess many pharmacological properties, including triterpenoid saponins with excellent antibacterial, antitumor, hepatoprotective, antihyperglycemic, antidyslipidemic, and insecticidal activities (Li, 1975; Upadhyay and Singh, 2012; Xu et al., 2018). In addition, soapberry pericarp has been traditionally used as a detergent because of its widespread ability to form stable soap-like foams in aqueous solution (Li, 1975; Upadhyay and Singh, 2012). To date, more than 70 triterpenoid saponin compounds have been identified in soapberry, which are mainly divided into four structural types, namely, oleanane, lupine, dammarane, and tirucullane types (Upadhyay and Singh, 2012; Xu et al., 2018). The majority of studies on soapberry saponins have focused on constituents and bioactivity, while there have been no reports regarding the biosynthesis of these physiologically active substances.

The biosynthesis of plant triterpenoid saponins is a complex process, and the whole pathway can be divided into three stages: isopentenyl diphosphate (IPP) and its isomer, dimethylallyl diphosphate (DMAPP), derived either from condensation of acetyl-CoA in the cytosolic mevalonic acid (MVA) pathway or from pyruvate and phosphoglyceraldehyde in the plastidial 2-C-methyl-D-erythritol 4-phosphate (MEP) pathway; IPP and DMAPP are catalyzed by the corresponding enzymes to produce 2,3-oxidosqualene; and 2,3-oxidosqualene is cyclized to a triterpenoid backbone by oxidosqualene cyclases (OSCs). These molecules are then chemically modified by cytochrome P450 monooxygenase (CYP450) and UDP-glycosyltransferase (UGT), resulting in the production of different types of triterpenoid saponins (Augustin et al., 2011; Yang et al., 2018; Zhao and Li, 2018; Xu et al., 2021). Several transcription factors (TFs) have been reported to be involved in the biosynthesis of triterpenoid saponins by regulating the expression of saponin biosynthetic genes, such as bHLH (Mertens et al., 2016), AP2/ERF (Deng et al., 2017), bZIP (Xu et al., 2019), and WRKY (Singh et al., 2017).

With the further development and improvement of multiomics technologies, the joint analysis of the metabolome and transcriptome has been widely conducted to determine the metabolic pathways and regulatory genes in numerous plant species, including *Solanum lycopersicum* (Li et al., 2020), *Capsicum annuum* (Liu et al., 2020), *Colchicum* spp., and *Gloriosa* spp. (Nett et al., 2020), *Citrullus lanatus* (Umer et al., 2020), *Actinidia chinensis* (Wang et al., 2021), and *Zea mays* (Zhang et al., 2021), as well as many trees, including *Ginkgo biloba* (Meng et al., 2019), *Dimocarpus longan* (Yi et al., 2020), and *Pyrus* spp. (Ni et al., 2020). However, there have been few studies involving integration of triterpenoid saponin metabolism and transcriptome data to explain the molecular regulatory mechanisms underpinning triterpenoid saponin biosynthesis in soapberry pericarps.

Here, we examined the triterpenoid saponin dynamics and the regulatory mechanisms underlying triterpenoid saponin biosynthesis in soapberry at eight pericarp growth stages. Through a combination of metabolite profiling and transcriptome analysis, saponin-specific clusters/modules were identified by Mfuzz clustering and weighted gene co-expression network analysis (WGCNA), and one TF–gene regulatory network controlling production and accumulation of triterpenoid saponin were constructed. In addition, we identified key TFs that modulate triterpenoid saponin biosynthesis metabolism by direct transcriptional regulation of their structural target genes. This study not only clarified the accumulation of saponins in soapberry pericarps during pericarp development, but also provided important insights into the regulation of triterpenoid saponin biosynthesis in soapberry.

## Results

### Identification of saponins in the soapberry pericarp at eight fruit growth stages

To investigate the constituents and dynamic changes of saponins in the developing soapberry pericarp, the total saponin content of soapberry pericarp was measured at eight fruit growth stages (Figure 1A; Figure 1B). Total saponins showed rapid accumulated from S1 (12.30%), moderate accumulated from S3 (19.04%), peaked at S4 (19.30%), and then decreased gradually, indicating that the saponin content changed dynamically during soapberry fruit growth. Metabolite profiling of pericarp was then performed at eight fruit growth stages by UHPLC-Q-Qrbitrap-MS. A total of 54 types of saponin (including 11 groups of isomers) were tentatively detected in all eight stages (Supplemental Table S1). Principal component analysis (PCA) confirmed that these saponins could be divided into two groups and exhibited clear separation among pericarp samples at different stages (Figure 1C). Based on their accumulation at different growth stages, the cluster dendrogram also distinguished two groups (Figure 1D), which was consistent with the PCA results. The saponins in cluster I (n = 9) accumulated preferentially at stage S1. Saponins in cluster II (n = 45) accumulated mainly at stage S2, S4, S5, or S8 (Figure 1D). These results suggested that the saponin profile shows a dynamically changing pattern during pericarp growth. In addition, saponins 19, 20, 22, 25, and 31 were the major saponins and accumulated specifically at stage S4. Saponins 11 and 12 were also the major saponins and accumulated preferentially at stage S8 (Supplemental Table S1; Figure 1D).

**Figure 1.**
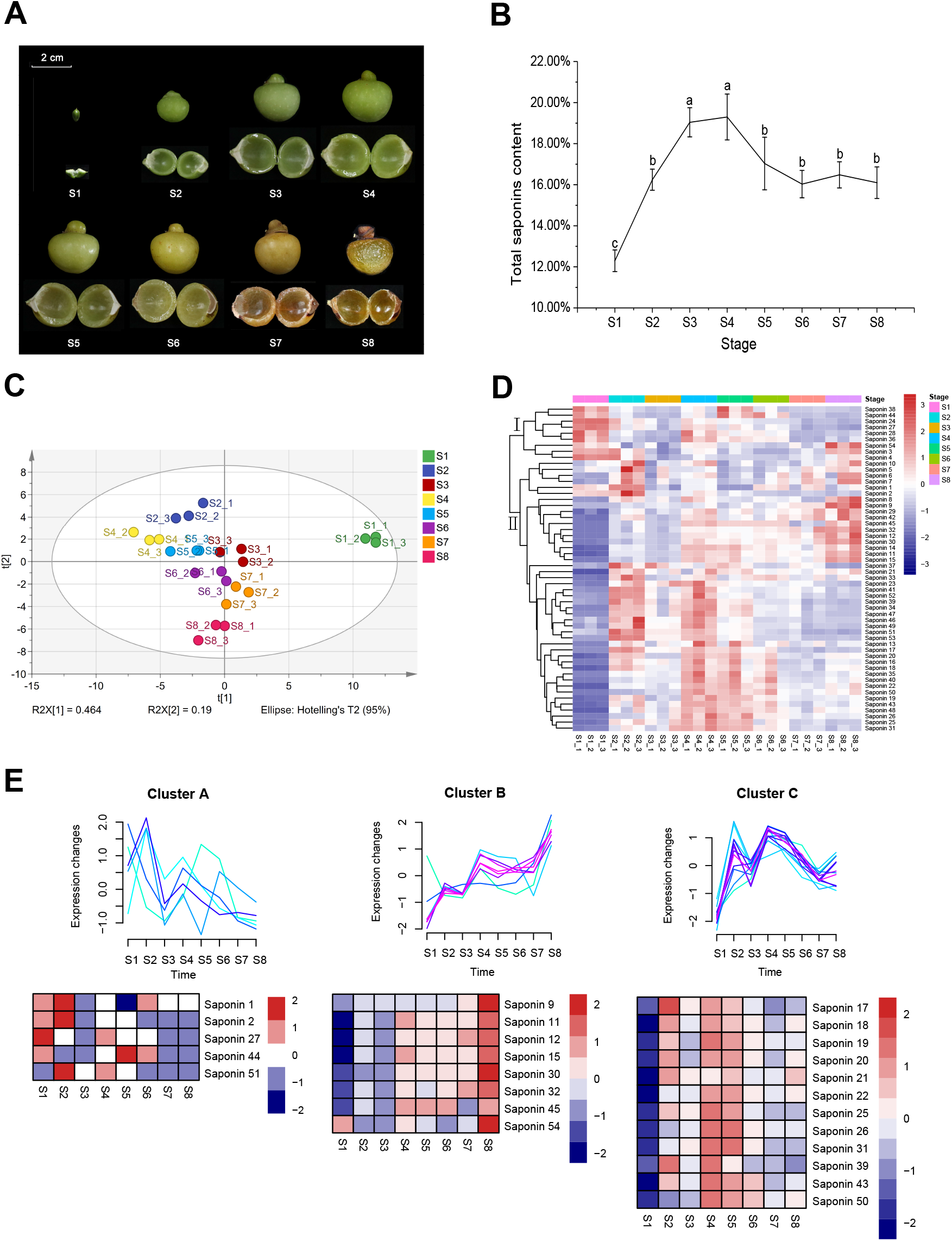
Images of soapberry fruits and dynamic changes of saponins in the pericarp at eight growth stages. (A) Images of soapberry fruits and pericarps at eight growth stages. (B) Dynamic changes of total saponin contents of soapberry pericarp at eight growth stages. Means ± SD (n=3). Means in different letters (a–b) are significantly different at *P*<0.05 by ANOVA on Duncan’s multiple range test. (C) Principal component analysis (PCA) of detected saponins. (D) Hierarchical clustering heat map of detected saponins. (e) Upper: Temporal expression patterns of DASs. All DASs were classified into three clusters based on their contents using the Mfuzz R package. (E) Lower: Heatmap of DASs accumulation level in cluster A, cluster B and cluster C.

Based on the pairwise comparison with |log_2_(fold change)| > 1, *p* < 0.05, and VIP (variable importance in project) > 1 as thresholds, a total of 25 differentially accumulated saponins (DASs) were identified (Supplemental Table S2). The 25 DASs were divided into three groups by Mfuzz clustering based on their accumulation levels (Ni et al., 2020) (Figure 1E; Supplemental Table S3). In cluster A, the saponins showed high accumulation at stage S1 or S2 and tapered off from stage S3 to S8. The content of saponins in cluster B increased continuously with fruit growth. For cluster C, the levels of saponin accumulation showed an uptrend trend, had high levels at stages S2, S4, and S5, and then decreased from S6 (Figure 1E).

### Transcriptome profiles of soapberry at eight growth stages

After removal of low-quality reads, ambiguous reads, and adapter sequences, each library had 31.97–86.81 M clean reads (Supplemental Table S4). These clean reads were mapped to the soapberry reference genome with average match ratios of 87.87%, and 23,783 genes predicted from the genome were found to be expressed in at least one sample with fragments per kilobase of transcript per million fragments mapped (FPKM) > 0. Furthermore, 9114 novel genes were also identified, which were not included in the reference genome. Twenty randomly selected genes were verified by qRT-PCR, which suggested that the relative expression levels were basically consistent with the RNA-Seq data (Supplemental Figure S1), thus supporting the accuracy and reliability of the transcriptomic profiling results. Through differential expression analysis between eight pericarp growth stages, a total of 22,002 DEGs were identified, and the DEG numbers increased gradually with fruit growth (Supplemental Figure S2). In addition, the number of downregulated genes was greater than that of upregulated genes (Supplemental Figure S2), suggesting that downregulated genes may play more important roles in soapberry saponin biosynthesis. GO enrichment analysis of these DEGs indicated that they were widely distributed in three functional groups. The genes related to *protein phosphorylation, transcription, DNA-templated,* and *carbohydrate metabolic process* were predominant in the *biological process* category. In the *cellular component* category, the genes were mostly involved in *endoplasmic reticulum membrane*, *vacuolar membrane*, and *vacuole*. In the *molecular function* category, *heme binding* was predominant followed by *iron ion binding* and *hydrolase activity* (Supplemental Figure S3).

Subsequently, the 22,002 DEGs were grouped into 12 clusters by Mfuzz clustering based on their expression levels (Supplemental Figure S4). Among them, the expression of clusters 2, 9, and 11 showed consistency with the accumulation patterns of DASs (clusters A, B, and C). DEGs from cluster 2 (2442 DEGs) showed similar expression tendencies, and their expression levels were highest at stage S1 and then decreased with fruit growth. Conversely, in cluster 11, the expression profiles of most DEGs (3816) were the opposite of those in cluster 2, showing an uptrend over time and peaking at stage S8. The DEGs (1560) in cluster 9 generally shared a trend of first increasing and then decreasing. Kyoto Encyclopedia of Genes and Genomes (KEGG) pathway enrichment analysis revealed that some of the DEGs in these three clusters were related to *terpenoid backbone biosynthesis* and *sesquiterpenoid and triterpenoid biosynthesis* (Supplemental Figure S5), suggesting that these DEGs play key roles in saponin biosynthesis in soapberry.

### Expression patterns of saponin biosynthesis-related genes in soapberry pericarp

To further explore the mechanism of saponin accumulation in soapberry pericarp, the expression patterns of genes possibly related to triterpenoid saponin biosynthesis were analyzed. In this study, we found 135 genes encoding the enzymes involved in the MVA pathway, MEP pathway, and saponin biosynthesis pathway (Figure 2; Supplemental Table S5). Among them, 49 genes were involved in putative backbone biosynthesis and some candidate genes were assumed to be responsible for backbone modification, including 41 CYP450s and 45 UGTs. In addition, phylogenetic analysis showed that most of the OSCs, CYP450s, and UGTs in our study had high degrees of identity with known examples (Supplemental Figure S6–S8). Most of the 135 candidate genes, such as 3-hydroxy-3-methylglutaryl-CoA reductase (HMGR), phosphomevalonate kinase (PMK), metholvalic-5-diphosphate decarboxylase (MVD), 4-cytidine diphosphate-2-C-methyl-D-erythritol kinase (CMK), 2-C-methyl-D-erythritol-2,4-cyclic phosphate synthetase (MDS), IPP isomerase (IDI), geranyl pyrophosphate synthetase (GPS), farnesyl pyrophosphate synthetase (FPS), squalene synthase (SS), squalene epoxidase (SE), β-amyrin synthase (β-AS), CYP450 and UGT, were highly expressed at the early stages of pericarp growth (stages S1–S4) (Figure 2). This was consistent with the trend that the contents of total saponins and most saponins were higher in the early pericarp growth stages (such as stages S2 and S4), indicating that saponins were mainly synthesized in the early stages of pericarp growth and we can further screen for genes closely related to saponin biosynthesis from these genes.

**Figure 2.**
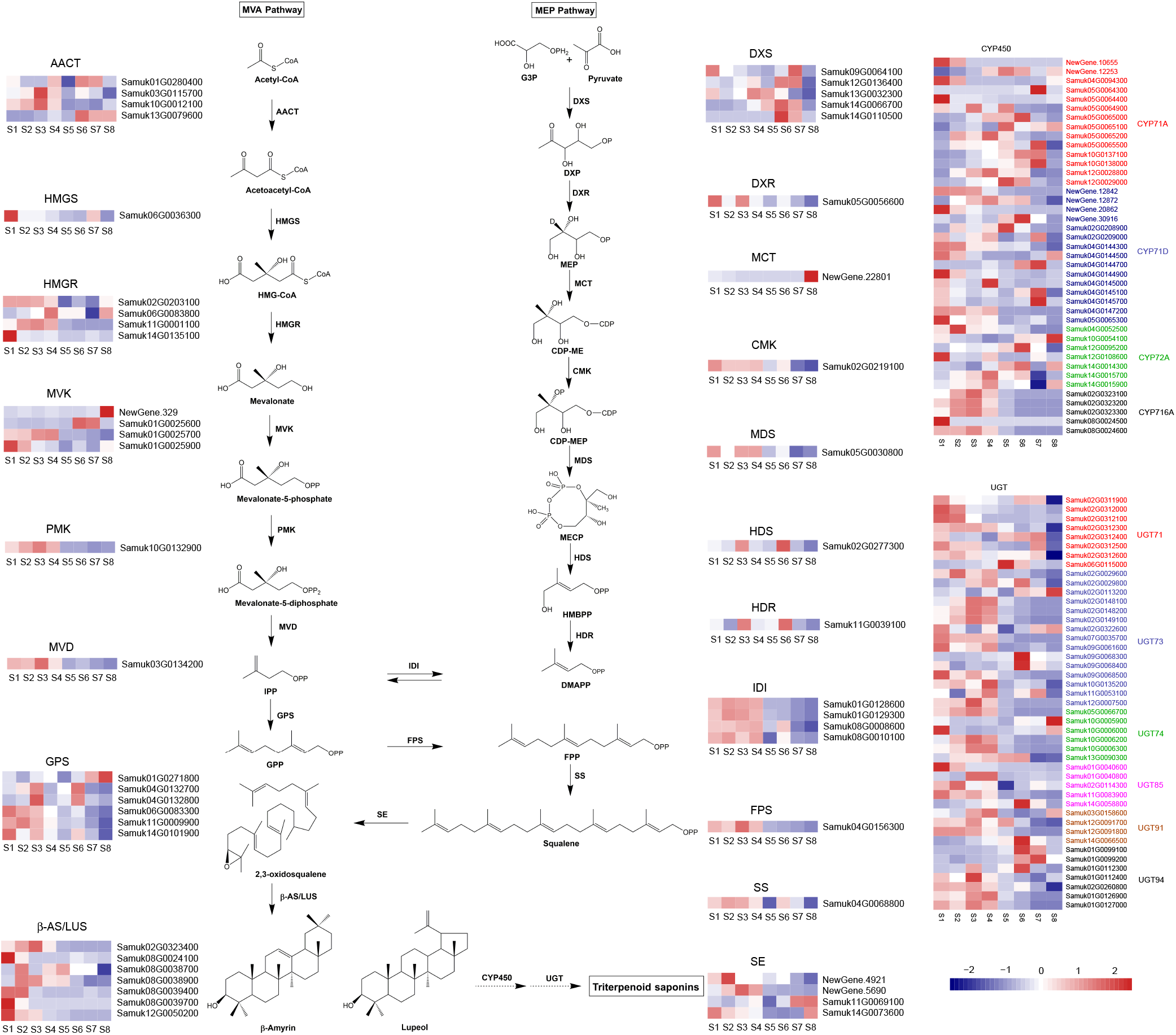
Proposed pathways for triterpenoid saponin biosynthesis in soapberry. Expression levels of genes encoding enzymes that catalyze each step of the triterpenoid saponin biosynthesis pathway are shown. The higher and lower expression level of genes are indicated in firebrick and navy, respectively.

### Expression patterns of TFs in soapberry pericarp

A total of 1785 TFs with different expression abundances were detected in the soapberry pericarp, which were classified into 54 families according to the Plant Transcription Factor Database (PlantTFDB) (Figure 3A). The MYB, NAC, WRKY, Nin-like, and ERF families of TFs accounted for the largest numbers of expressed TFs, with 201, 181, 180, 164, and 110 members, respectively. Dynamic analysis showed that most TF families had high expression levels at stages S1–S4. Meanwhile, some TF families had higher FPKM levels than other TF families, such as ARF, bZIP, CAMTA, DBB, Dof, MYB, and WRKY (Figure 3A). In addition, the expression patterns of TFs from clusters a, d, and f by Mfuzz clustering were consistent with the accumulation of DASs (clusters A, B, C), including 130 MYBs, 114 WRKYs, 67 ERFs, 57 bHLHs, and 36 bZIPs, suggesting that these TFs may play important roles in regulating saponin biosynthesis in soapberry (Figure 3B).

**Figure 3.**
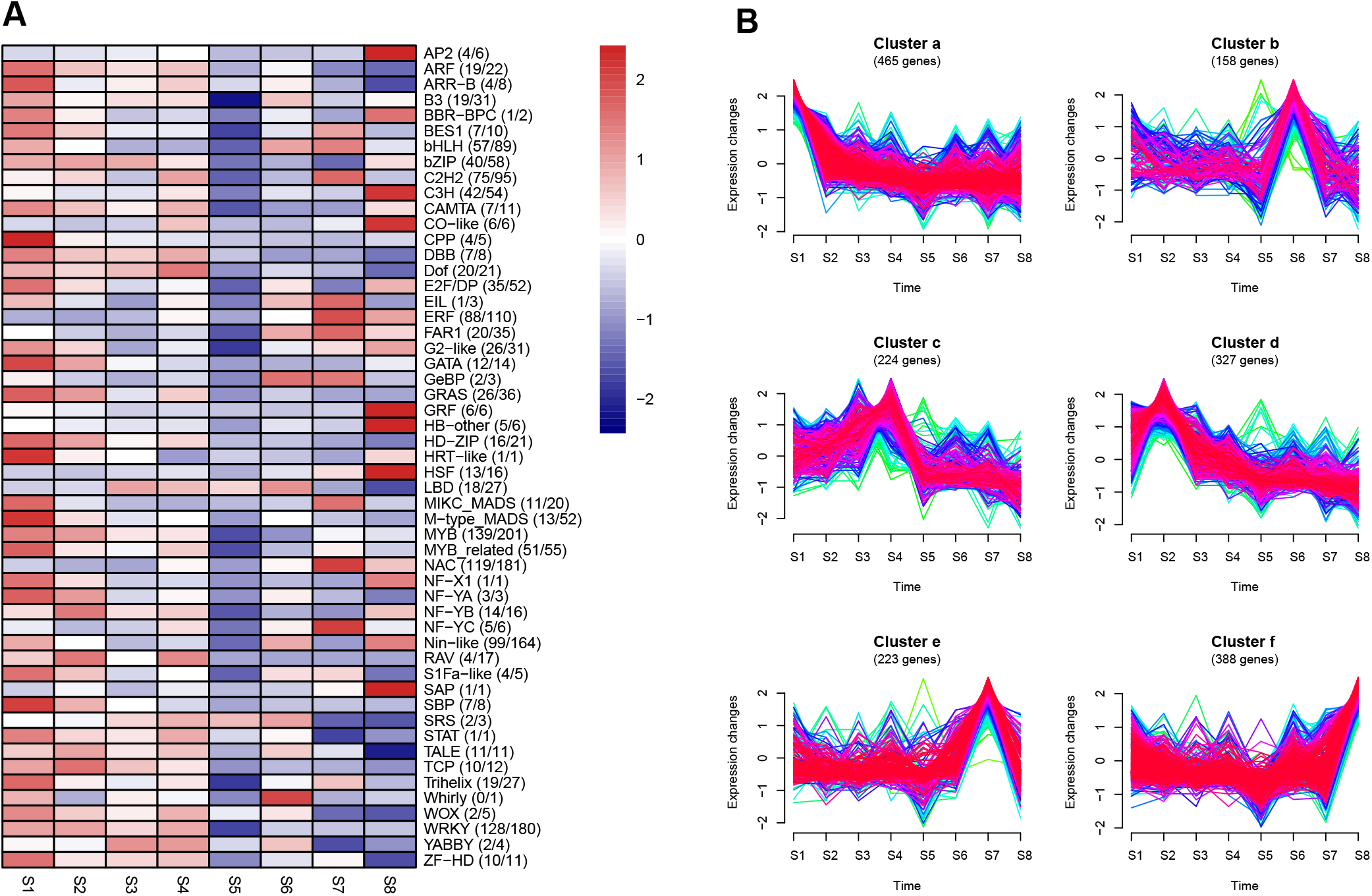
Expression dynamics of TF families. (A) The colour shows the total FPKM of all TFs of a particular TF family. The first number in parentheses is the number of DEGs in a TF family, and the second number refers to the total number of genes in that TF family as annotated in PlantTFDB. (B) Expression profiles of all TFs by Mfuzz clustering analysis.

### Co-expression network analyses identified saponin-related genes

To obtain further insight into the triterpenoid saponin biosynthesis in soapberry, WGCNA was performed to investigate the co-expression networks of all expressed genes. After deletion and outlier values were eliminated, 18,717 genes remained. These genes were clustered into 24 major tree branches, each of which represented a module (labeled with different colors; the grey module contains the remaining uncorrelated genes) (Figure 4A; Supplemental Table S6). Analysis of module–saponin relationships showed that the blue and greenyellow modules were significantly correlated with most saponins, with high correlation coefficients (Figure 4B). The blue module with 5211 genes was highly positively correlated with saponins 1, 2, and saponin 27, and highly negatively correlated with total saponins and 15 other saponins (including saponins 19, 20, and 22). The greenyellow module contained 3090 genes and showed a significant association with saponins 9, 12, 30, 32, and 54. Therefore, we selected these two modules (containing 8301 genes) positively or negatively correlated with saponin biosynthesis for further study. Genes preferentially expressed in the early stages of fruit growth (stages S1–S4) were mainly accumulated in the blue module (Figure 4C), while most of the genes highly expressed in the fully developed and mature stage (stage S8) were located in the greenyellow module (Figure 4D). Although both of these modules showed strong correlations with saponins, the genes in the two modules had different expression patterns, suggesting that the genes in these two modules may be involved in different functions in the biosynthesis of saponins. In addition, the patterns of changes in gene expression in the two modules were consistent with the results of Mfuzz clustering analysis, where 2442 genes (cluster 2) were found to be highly expressed in the early stages of fruit growth (stage S1) and 3816 genes (cluster 11) were highly expressed in stage S8 (Figure 4D). In addition, most of the genes in these two modules were also identified in the selected Mfuzz clusters. Thus, both WGCNA and Mfuzz clustering have identified two clusters/modules consistent with saponin accumulation patterns despite the different algorithms.

**Figure 4.**
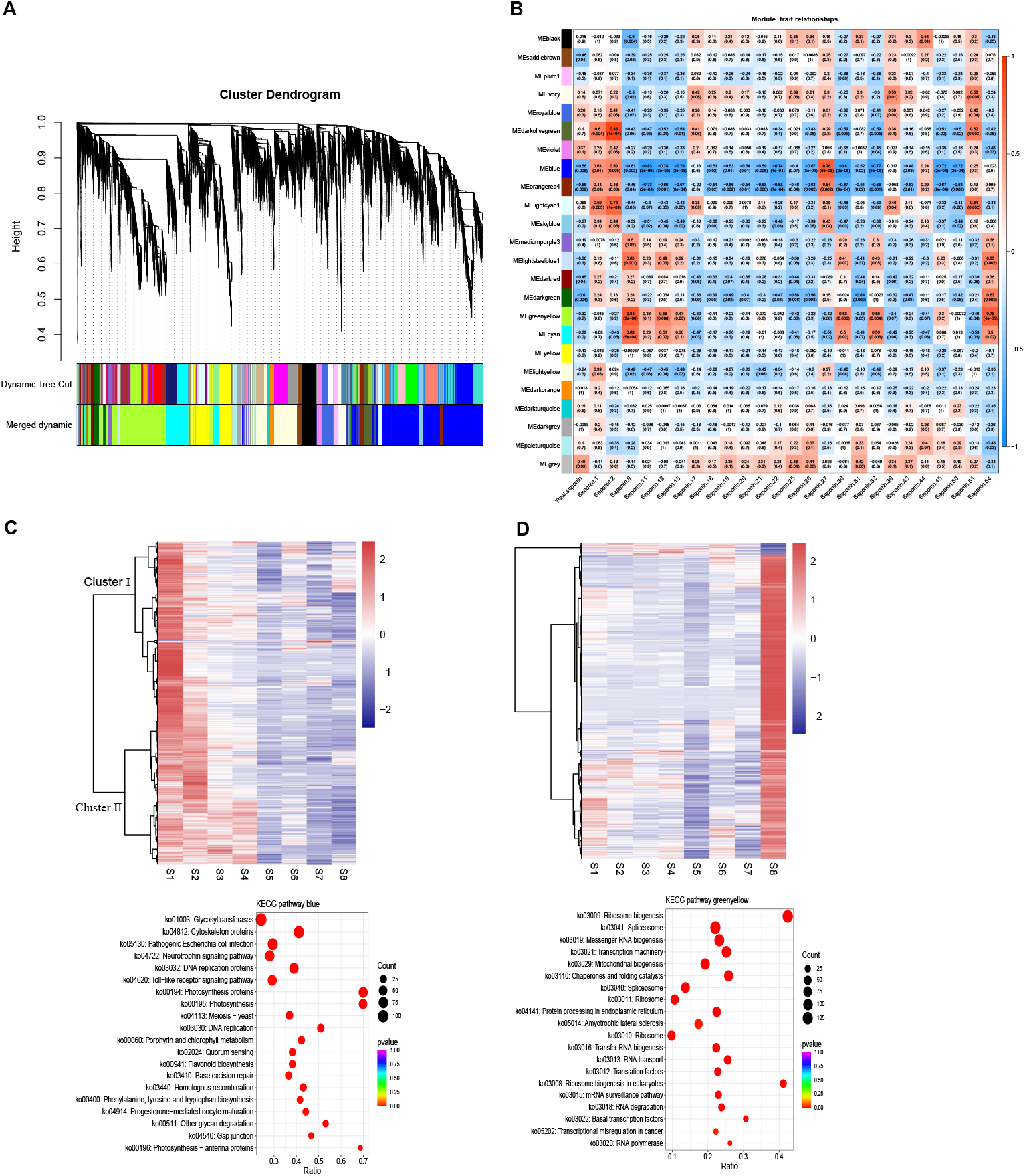
WGCNA of all expressed genes. (A) Hierarchical clustering tree (cluster dendrogram) showing 24 modules of co-expressed genes by WGCNA. Each leaf of tree corresponds to one gene. The major tree branches constitute 24 modules, labeled with different colors. (B) Module-saponin relationship. Each row represents a module. Each column represents a specific saponin (detailed information reference Table S4). The value in each cell at the row-column intersection represents the correlation coefficient between the module and the saponin and is displayed according to the color scale on the right. The value in parentheses in each cell represents the *P* value. (C) Upper: heat map of genes in the blue module; Lower: the KEGG enrichment analysis of the genes in the blue module showed only the top 20 pathways with the most significant enrichment. (D) Upper: heat map of genes in the greenyellow module; Lower: the KEGG enrichment analysis of the genes in the greenyellow module showed only the top 20 pathways with the most significant enrichment.

Further, the results of KEGG analysis results showed that the genes in the blue module were primarily related to *glycosyltransferases*, *photosynthesis*, *photosynthesis proteins*, and *cytoskeleton proteins* (Figure 4C), which were strongly correlated with saponins, suggesting that glycosyltransferases and photosynthetic metabolites may play important roles in saponin biosynthesis. This is similar to the conclusion that the photosynthetic metabolites (glucose and fructose) contribute to the synthesis of ginsenosides or movement into ginseng roots reported by schramek et al (Schramek et al., 2014). In addition, 17 saponin biosynthetic structural genes, including *HMGR, DXS, GPS* and *β-AS,* were enriched in *terpenoid backbone biosynthesis* and *sesquiterpenoid and triterpenoid biosynthesis*. Meanwhile, another 27 saponin biosynthesis candidate genes, including CYP450s and UGTs, were also found in the blue module. These results further verified that the co-expressed genes in this module are involved the accumulation of saponins in the soapberry pericarp. In the greenyellow module, the genes were significantly enriched for *ribosome biogenesis*, *messenger RNA biogenesis*, *transcription machinery*, and *basal transcription factors* (Figure 4D), suggesting that the genes in this module may regulate the biosynthesis of saponins.

### Candidate hub genes related to saponin biosynthesis

Module hub genes are usually considered as representatives of given modules in biological networks. In the blue module, 46 genes showed strong correlations (edge weight ≥ 0.35) (Supplemental Table S7; Supplemental Figure S9A). Among them, kinesin family member 4/21/27 protein (*KIF4_21_27, Samuk02G0147800),* putative protein kinase RLK-Pelle-LRR-XIIIb family protein (*RPLX1, Samuk12G0107600),* INO80 complex subunit B protein (*INO80B, Samuk13G0014100), Samuk05G0034700* (unknown protein), cyclin D3 (*CYCD3*, *Samuk02G0146500*), LRR receptor-like serine/threonine-protein kinase ERECTA (*ER*, *Samuk06G0040200*), putative X8 domain-containing protein (*X8DC1*, *Samuk05G0140500*), and *SmCYP71D-3 (NewGene.20862)* had the highest degree of connectivity in the network, suggesting that these genes may play important roles in saponin biosynthesis. The network diagram of the greenyellow module is composed of 42 co-expressed genes (edge weight ≥ 0.45), in which genes such as the BES1 family protein (*SmBES1-7, Samuk03G0151400),* protease Do-like 9 (*PD9*, *Samuk10G0098700),* DNA-directed RNA polymerase (*RPO41, Samuk09G0117400*), transcription elongation regulator 1 (*CA150*, *Samuk07G0143300*), and MYB family protein (*SmMYB57*, *Samuk08G0065200*) have the highest degree of connectivity (Supplemental Table S8; Supplemental Figure S9B), suggesting that these genes may also participate in saponin biosynthesis. However, this requires further verification.

### Generation of the saponin biosynthesis regulatory network

To construct the regulatory network associated with saponin biosynthesis, we screened the structural genes involved in the saponin biosynthesis pathway identified in the blue and greenyellow modules. We identified 44 structural genes associated with saponin biosynthesis in the blue module, the expression levels of which were strongly correlated with the accumulation of saponins (Supplemental Table S9). By associating the expression patterns and the potential binding affinity for the promoters of these saponin biosynthesis-related genes, we identified 40 TFs, including ERF, TCP, SBP, Trihelix, CPP, bHLH, and WRKY—the expression levels of which were strongly correlated with the 30 saponin biosynthesis-associated genes in the blue module—and generated a correlated network (Supplemental Table S10; Figure 5). These observations indicated that these TFs correspond to the putative regulators controlling saponin biosynthesis in soapberry pericarp. As only four saponin biosynthesis candidate genes were identified in the greenyellow module (Supplemental Table S11), no eligible structural TF–gene pairs were detected. In addition, the results of qRT-PCR showed that the expression levels of multiple TFs in the blue module exhibited strong correlations (Pearson correlation coefficient, r > 0.6) with structural genes—for example, *SmHMGR4, SmCYP71D-3, SmCYP716A-5,* and *SmUGT91A-2—*thus further verifying the reliability of the regulatory network(Figure 6).

**Figure 5.**
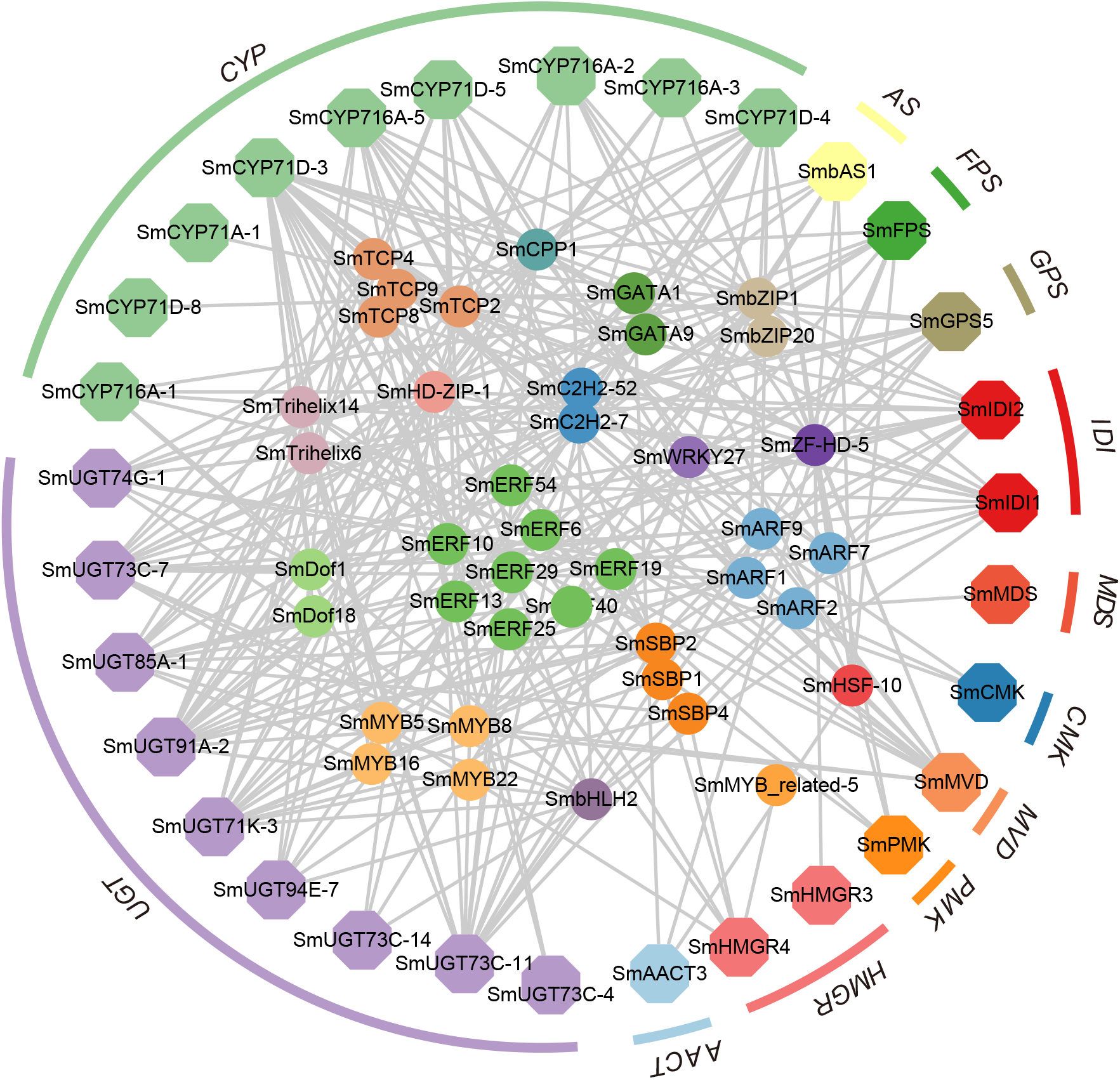
The regulatory network of triterpenoid saponin biosynthesis in soapberry. Hexagon with different colors indicate different families of structural genes associated with saponin biosynthesis in the blue module. Circles with different colors indicate different families of TFs characterize in the same module which transcripts are highly correlated with expression of structural genes.

**Figure 6.**
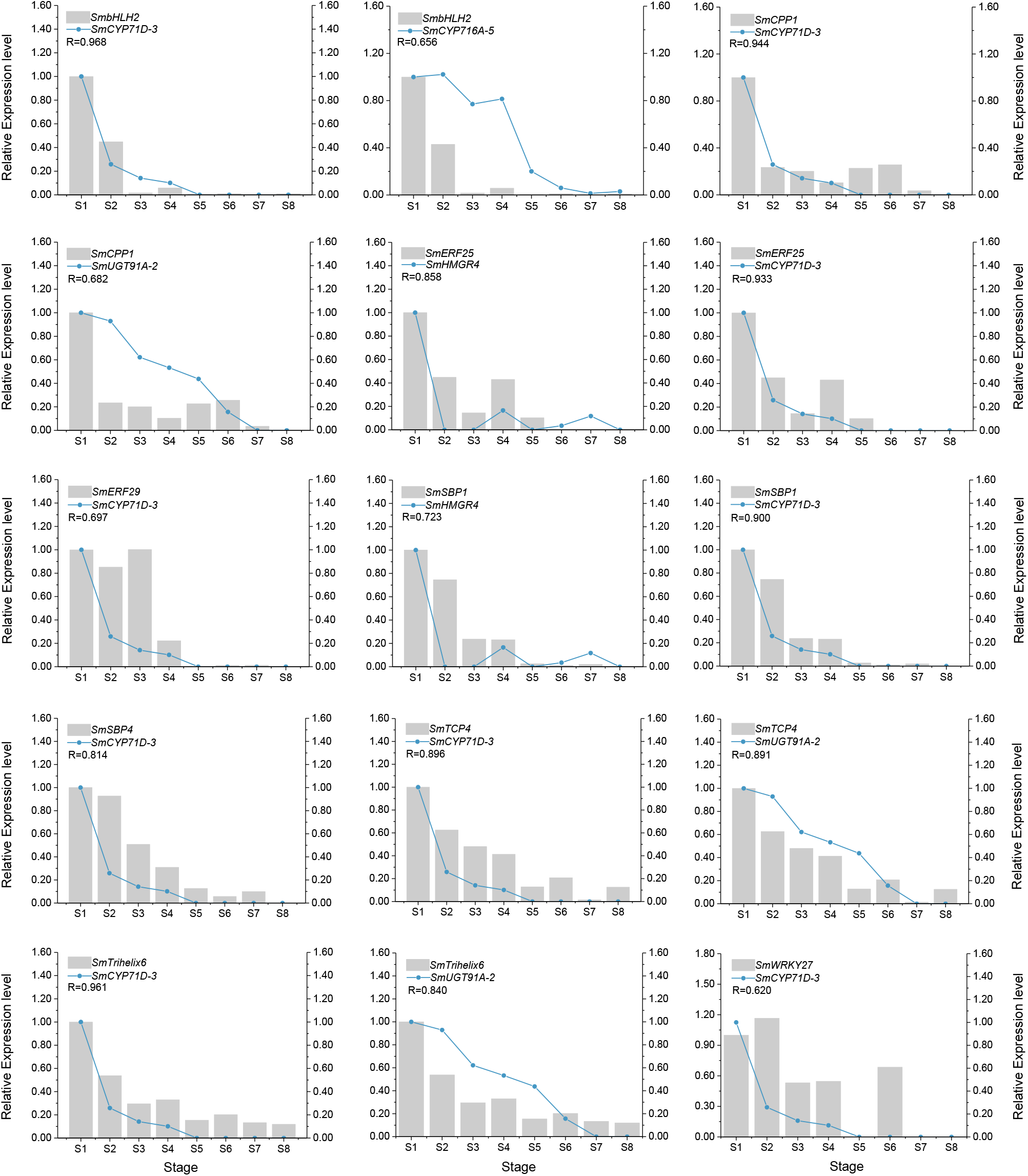
Expression correlation of TFs and their predicted target genes. The bars and lines indicate the relative expression level of TFs and their corresponding target genes in the eight growth stages of soapberry pericarps. The Y-axis on the left and right represents the abundance of the TFs and their target genes, respectively. *SmACT* was used for normalizing the relative expression of these encoding genes. The expression level of the genes in the stage S1 were set as 1.0. Relative expression level was calculated using the 2^-ΔΔ*C*t^ method. Data indicate the mean values of three biological replications.

### Identification of key TFs regulating saponin biosynthesis

As structural modification of the triterpenoid backbone is one of the most important steps in the biosynthesis of triterpenoid saponins, we focused our analysis on the regulatory network of CYP450 genes responsible for oxidizing the backbones created by OSCs into sapogenins. There were nine CYP450s associated with saponin biosynthesis in the TF–structural gene regulatory network in the blue module (Figure 5). To identify the key TFs modulating activation of the candidate genes encoding CYP450s, we focused on the regulation of *SmCYP71D-3* because this candidate gene exhibited high levels of transcript accumulation as well as expression specificity and was identified as a DEG and a hub gene in the blue module (Supplemental Figure S10). This regulatory network analysis suggested that *SmCYP71D-3* is highly correlated with 29 TFs, including the ERF, TCP, SBP, bHLH, CPP, Trihelix, and WRKY families (Supplemental Table S12; Supplemental Figure S10). We randomly selected nine of these TFs as candidates for further study, namely, SmbHLH2, SmCPP1, SmERF25, SmERF29, SmSBP1, SmSBP4, SmTCP4, SmTrihelix6, and SmWRKY27, which showed the strongly correlated expression, high expression levels, and the ability to bind to the promoter of *SmCYP71D-3* (Figure 7A; Supplemental Figure S10). Further, we conducted yeast one-hybrid (Y1H) assays to investigate direct binding of these nine TFs to the *SmCYP71D-3* promoter. The fragment containing the core motifs of the *SmCYP71D-3* promoter was cloned into the upstream of the Aureobasidin (AbA) resistance gene and transformed into the Y1H Gold strain as bait. In addition, the basal activity of the Y1H Gold strain transformed with pAbAi-*SmCYP71D-3* was successfully inhibited in the presence of 50 ng/mL AbA (Figure 7A). We expressed these TFs fused to the GAL4 AD in the Y1H system to challenge them with the *SmCYP71D-3* promoter fused to the *AbAr* reporter. Among the nine TFs, only the yeast transformed with SmbHLH2, SmTCP4, and SmWRKY27 grew well in the selection medium, suggesting that these three TFs could induce *SmCYP71D-3* promoter activity (Figure 7A). Consistent with these observations, electrophoretic mobility shift assays (EMSAs) suggested that SmbHLH2, SmTCP4, and SmWRKY27 bound to the DNA probes containing the corresponding TF binding sites present in the promoter region of *SmCYP71D-3* (Figure 7B). In conclusion, these results support the hypothesis that SmbHLH2, SmTCP4, and SmWRKY27 may play important roles in the accumulation of saponin via direct regulation of the transcription of *SmCYP71D-3* in soapberry pericarp.

**Figure 7.**
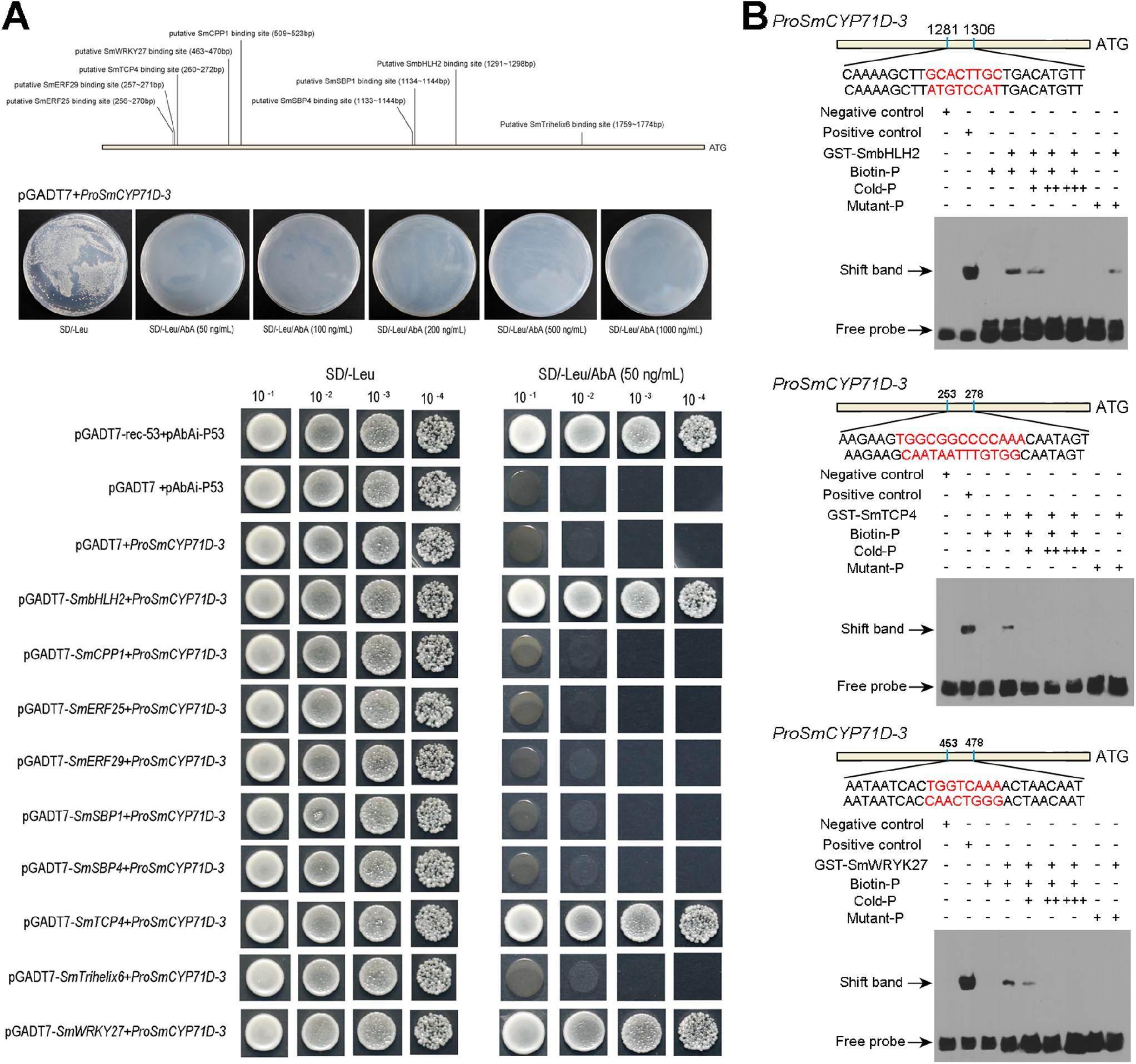
Identification of SmbHLH2, SmTCP4, and SmWRKY27 as key TFs regulating *SmCYP71D-3* in soapberry. (A) Upper: schematic illustration of the potential recognition elements of the nine TFs in the 2 kb promoter region of *SmCYP71D-3*. (A) Middle: no basal activities of *ProSmCYP71D-3* were observed in yeast grown on SD medium lacking Leu in the presence of 50, 100, 200, 500, and 1000 ng/mL^-1^ AbA (SD/-Leu/AbA). (A) Lower: yeast growth assays after Y1H reporter strains were transformed with effector containing the ORF of different TFs or empty (negative control). Interaction was determined based on the ability of transformed yeast to grow on SD medium lacking Leu in the presence of 50 ng/mL^-1^ (SD/-Leu/AbA). (B) SmbHLH2, SmTCP4, and SmWRKY27 binding to the promoter of *SmCYP71D-3*. The WT probe containing the corresponding TF binding site was biotin-labeled. Competition for TF binding was performed with cold probes. The symbols - and + represent absence or presence of the probes and GST-tagged TF protein. The core binding sequences are highlighted red; The upper sequence is the prediction TF binding site sequence, and the lower sequence is the mutation probe sequence.

## Discussion

Soapberry pericarp has been used as medicine and as an additive for cosmetics due to its abundant triterpenoid saponins with several bioactivities and abilities as foaming agents. However, very little is known about saponin accumulation or the molecular mechanism of saponin biosynthesis in soapberry pericarp. In this study, the differences in saponins at different growth stages of soapberry pericarp were analyzed qualitatively and quantitatively analyzed by UHPLC-Q-Qrbitrap-MS (saponin targeted). Metabolite profiling tentatively identified 54 saponins, significantly increasing our understanding of the saponins in soapberry pericarp. Saponin-producing plants usually accumulate saponins as part of their normal development. Meanwhile, saponin accumulation is also affected by many environmental factors, including nutrient and water availability, light irradiation, diseases, and pests (Augustin et al., 2011). Ndamba et al. (1994) suggested that maximal saponin accumulation in the early stages of *Phytolacca dodecandra* berry growth is to prevent fruit loss and ensure seed maturation. We obtained similar results indicating that the most saponins in soapberry pericarp accumulated rapidly in stage S2 and reached the highest level in stage S4 (Figure 1), accompanying by two fruit loss stages (Zhao et al., 2019). Meanwhile, the precipitation and temperature in the orchard were highest at stage S4 (Zhao et al., 2019). It was inferred that variation in saponin levels in the soapberry pericarp may be related to protection and responses to environmental factors.

Plant triterpenoid saponins have a wide variety of complex structures (Augustin et al., 2011). Although they have a common precursor synthesis pathway, the post-modification process from 2,3-oxidosqualene to the formation of the final triterpenoid saponin has various rules in different plants (Haralampidis et al., 2002; Augustin et al., 2011). This process mainly involves three enzymes, namely, OSCs, CYP450s, and UGTs. Cyclization of 2,3-oxidized squalene is considered the first committed step in the triterpenoid saponin biosynthesis. Then, the basal sapogenin backbone formed by OSCs usually undergoes hydroxyl, ketone, aldehyde and carboxyl modification by CYP450s before glycosylation. Finally, the saccharide side chains are introduced into the sapogenin backbone by UGTs to finalize the biosynthesis of saponin. A total of 135 saponin biosynthesis candidate genes were annotated in the persent study, including 7 OSCs, 41 CYP450s, and 45 UGTs. Among them, four *β-ASs (SmbAS1, SmbAS3, SmbAS4,* and *SmbAS6*) showed identity to *Betula platyphylla* var. *japonica BPY* (Zhang et al., 2003), *Lotus japonicus LjAMY1* (Sawai et al., 2006), and *Glycyrrhiza glabra GgbAS1* (Hayashi et al., 2001), suggesting that they may encode β-amyrin synthase catalyzing conversion of 2,3-oxidosqualene to β-amyrin (Supplemental Figure S6). Another candidate *OSC*, *SmLUS,* showed a high degree of identity to *B. platyphylla* var. *japonica BPW* (Zhang et al., 2003), *L. japonicus OSC3* (Sawai et al., 2006), and *G. glabra GgLUS1* (Hayashi et al., 2004), suggesting that this *OSC* may be responsible for the biosynthesis of lupane-type triterpenoid saponin in soapberry pericarp. Furthermore, the CYP450 enzymes, which catalyze the oxidation of β-amyrin or lupeol, especially at C-3, C-23, and C-28, are required for the biosynthesis of the main triterpenoid saponins in soapberry pericarp (Xu et al., 2018). CYP716As have been shown to be mainly responsible for catalyzing sequential three-step oxidation of the α-amyrin/β-amyrin/lupeol backbone to form hydroxyl, aldehyde and carboxyl moieties at the C-28 position (Ghosh, 2017; Miettinen et al., 2017; Xu et al., 2021). In this study, two CYP716As, *SmCYP716A-2* and *SmCYP716A-3*, showed some similarity to *Chenopodium quinoa CqCYP716A78* (Fiallos-Jurado et al., 2016) and *Aquilegia coerulea AcCYP716A110* (Fiallos-Jurado et al., 2016) which catalyze β-amyrin to oleanolic acid, suggesting that they have the same catalytic activities in soapberry pericarp (Supplemental Figure S7). We also found that three candidate CYP72As (*SmCYP72A-1*, *SmCYP72A-5*, and *SmCYP72A-7*) showed identity to *Kalopanax septemlobus KsCYP72A397*, (Han et al., 2017) which catalyzes the hydroxylation of oleanolic acid at C-23, suggesting that these CYP72As may catalyze oleanolic acid to hederagenin in soapberry (Supplemental Figure S7). In addition, four CYP71Ds, *SmCYP71D-1, SmCYP71D-2, SmCYP71D-3,* and *SmCYP71D-4,* showed identity to *LjCYP71D353,* which catalyzes the C-20 hydroxylation of dihydro-lupeol and C-28 oxidation of 20-hydroxy-lupeol in *L. japonicus* (Krokida et al., 2013), suggesting that these candidate CYP71Ds may participate in the biosynthesis of lupane-type triterpenoid saponins in soapberry (Supplemental Figure S7). UGTs participating in saponin biosynthesis belonging to the UGT 71, 73, 74, 85, and 91 clans were identified previously (Rahimi et al., 2019). The UGT73 clan is the best candidate group for oleanane-type triterpenoid saponin biosynthesis (Erthmann et al., 2018). We identified three genes (*SmUGT73C-9, SmUGT73C-10,* and *SmUGT73C-11*) belonging to the UGT73 clan that showed high degrees of identity to *Panax japonicus* var. *bipinnatifidus PjOAGT* and *Panax zingiberensis PzOAGT1, PzOAGT2,* and *PzOAGT3* (Supplemental Figure S8), which transfer glucuronic acid at C-3 of oleanolic acid to form oleanolic acid 3-O-β-glucuronide (Tang et al., 2019), suggesting that these genes may catalyze the glucuronosylation of the C3-hydroxyl group for the biosynthesis of triterpenoid saponins in soapberry. In addition, four candidate UGT74s (*SmUGT74G-1, SmUGT74G-2, SmUGT74G-3,* and *SmUGT74G-4*) two candidate UGT94s (*SmUGT94E-1* and *SmUGT94E-2*), and six candidate UGT71s (*SmUGT71D-1*, *SmUGT71D-2*, *SmUGT71D-3*, *SmUGT71D-4*, *SmUGT71K-1*, and *SmUGT71K-4*) may be responsible for the glycosylation of tetracyclic sapogenins in soapberry (Dai et al., 2015; Wang et al., 2015; Wei et al., 2015; Lu et al., 2017; Lu et al., 2017). Further studies are needed to determine the functions of these candidate genes in the triterpenoid saponin biosynthesis pathway, including the key intermediates, in soapberry.

Combining these transcriptomic and metabolomic resources allowed the identification of key structural and regulatory genes involved in triterpenoid saponin biosynthesis in soapberry. This study not only validated some previously reported metabolic regulatory network information, but also identified novel TFs that regulate triterpenoid saponin biosynthesis. Previous studies have identified two bHLH family TFs, TSAR1 and TSAR2, from *Medicago truncatula,* that could instigate the expression of *HMGR1, β-AS, CYP93E2, CYP72A61v2, UGT73K1,* and *UGT73F3* as well as *MAKIBISHI1, β-AS, CYP716A12, CYP72A68v2,* and *UGT73F3*, resulting in corresponding accumulation of nonhemolytic and hemolytic soyasaponins, respectively (Mertens et al., 2016). In this study, we verified the relatively high correlation of expression (r=0.96) between *SmbHLH2* and *SmCYP71D-3,* which suggested the reliability of our analysis to identify regulators that control saponin biosynthesis. Y1H assay and EMSA also confirmed the transcriptional regulatory action of SmbHLH2 on *SmCYP71D-3*. In addition, the correlations in expression between SmCPP1, SmERF25, SmERF29, SmSBP1, SmSBP4, SmTCP4, SmTrihelix6, SmWRKY27, and *SmCYP71D-3* were also high (*r*>0.80). Considering the much higher expression levels of these TFs throughout pericarp development, we proposed that these TFs may be key regulators involved in modulating triterpenoid saponin accumulation. qRT-PCR results also showed that the expression pattern of *SmCYP71D-3* was similar to those of SmCPP1, SmERF25, SmERF29, SmSBP1, SmSBP4, SmTCP4, SmTrihelix6, and SmWRKY27 (Figure 6). However, Y1H assay and EMSA results showed that only SmTCP4 and SmWRKY27 would be able to bind to *SmCYP71D-3* via the predicted *cis*-motif elements, indicating that the results of bioinformatics prediction included certain false positives. The physiological function of *SmCYP71D-3* and corresponding TFs in saponin metabolism and the transcriptional regulation of these TFs on *SmCYP71D-3* by these TFs require further study. Further investigations focusing on the functions of other structural and regulatory genes would help to elucidate the molecular regulatory mechanisms of soapberry pericarp saponin biosynthesis.

## Materials and methods

### Plant materials

For this experiment, three 10-year-old soapberry superior trees were selected for sampling. The details of these plant materials were described in our previous report (Xu et al., 2021). The plant materials were cultivated at soapberry national forest germplasm banks in Jianning County, Fujian Province, China under natural conditions. Fruit samples were collected at eight growth stages between June and November 2018: early ovary development (stage S1), approximately 15 days after pollination (DAP); 30% of the largest fruit size (stage S2), 45 DAP; 70% of the largest fruit size (stage S3), 75 DAP; 80% of the largest fruit size (stage S4), 90 DAP; 90% of the largest fruit size (stage S5), 105 DAP; beginning of maturity (stage S6), 120 DAP; great change in pericarp (stage S7), 135 DAP; fully developed and mature (stage S8), 150 DAP. After picking the fruit, we immediately separated the pericarp from the seed. A portion of each pericarp sample was transferred to liquid nitrogen, and finally stored at −80 °C for RNA extraction. The other portion of each sample was dried to a constant weight in an oven at 40°C and then ground into uniform powder with a ball mill (MM400; Retsch, Arzberg, Germany) for saponin determination.

### Total saponin measurement and saponin metabolite profiling

The total saponin content was measured by the vanillin-glacial acetic acid method, as described by Sun et al. (2017). Absorbance was measured with a UV-Vis spectrometer (Lambda 35; PerkinElmer, Wellesley, MA, USA) at 547 nm. All experiments were performed in three biological replicates and three technical replicates.

The relative quantities of saponin metabolites in soapberry pericarp samples were analyzed by ultra-high-performance liquid chromatography with a Q Exactive Hybrid Quadrupole Orbitrap High-Resolution Accurate Mass Spectrometry (UHPLC-Q-Qrbitrap-MS) system by Shanghai Profleader Biotech Co., Ltd., Shanghai, China) (refer to Supplemental Method S1 for details regarding the protocol).

### RNA extraction, library construction, and sequencing

Total RNA extraction was performed using a FinePure Plant RNA Kit (Genfine Biotech Co. Ltd., Beijing, China) and the qualities were assessed using a Bioanalyzer 2100 (Agilent, Palo Alto, CA, USA). A cDNA library was constructed using a NEBNext® Ultra™ RNA Library Prep Kit for Illumina® (New England Biolabs, Ipswich, MA, USA) in accordance with the manufacturer’s protocol. A Qubit fluorometer® (Thermo Fisher Scientific, Waltham, MA, USA) and Qseq100 DNA Analyzer (Bioptic Inc., San Francisco, CA, USA) were used to determine the quality of the library based on the criteria of DNA concentration was > 1.0 ng/μL and the DNA length ~ 400 bp. The DNA molar concentration of the library was quantified with the KAPA Library Quantification Kit Illumina® Platform (KAPA Biosystems, Woburn, MA, USA). Libraries were sequenced using an Illumina HiSeq® X Ten System with paired-end reads by Beijing Yuanyi Biotechnology Co., Ltd. (Beijing, China).

### Analysis of RNA-Seq data

First, the adapter sequences, reads containing Ns (uncertain base information), and low-quality sequence reads were removed from the raw reads to produce clean reads. These clean reads were then aligned to the soapberry reference genome (Unpublished) sequence to acquire the genes using HISAT2 (Kim et al., 2015) and StringTie software (Pertea et al., 2015). The transcripts consistent with the reference genome were called known genes, and other new transcripts were called novel genes (prefixed with “NewGene”). The annotation information of genes was obtained from the soapberry genome annotation project. In addition, we also aligned the genes to annotated protein sequences of two species containing triterpenoid saponin, *Medicago truncatula* (http://www.medicagogenome.org/) and *Glycine max* (https://www.soybase.org/sbt/) (BLASTX; e-value ≤ 1e-5). To analyze candidate TFs in soapberry, nucleotide sequences were compared with the sequences in the Plant Transcription Factor Database (PlantTFDB) (http://planttfdb.gao-lab.org/index.php) using the default parameters (Tian et al., 2019).

Gene expression levels were calculated and normalized to fragments per kilobase of transcript per million fragments mapped (FPKM) (Trapnell et al., 2010). The differentially expressed genes (DEGs) between the samples were identified with DESeq2 (1.20.0) (Anders and Huber, 2010; Love et al., 2014). Genes with an Benjamini-Hochberg-adjusted *p*-values (*Q*-values) < 0.05 and |log_2_(fold change)|>1 were considered to be differentially expressed genes (DEGs). The identified DEGs were further subjected to enrichment analysis through Gene Ontology (GO) annotation and Kyoto Encyclopedia of Genes and Genomes (KEGG) pathway analysis using the clusterProfiler R package.

The deduced amino acid sequences of the *OSC*, *CYP450*, and *UGT* genes of soapberry in this study and those of other plants obtained from GenBank were used for phylogenetic analysis. Multiple sequence alignments were generated using Clustal W program. Phylogenetic analysis was carried out using the Neighbor-joining method with MEGA-X software program. A bootstrap of 1000 replications was used to estimate the strength of nodes in the tree.

### WGCNA and gene network visualization

The gene co-expression networks were constructed by the WGCNA method implemented in R software based on all expressed genes in our study (Langfelder and Horvath, 2008). The co-expression modules were constructed using the automatic network construction function (blockwiseModules) with default parameters, except for the soft thresholding power of 9, mergeCutHeight of 0.25, deepSplit of 2, and minModuleSize of 30. FPKM values were normalized, and an adjacency matrix was constructed. The resulting adjacency matrix was converted to a topological overlap matrix (TOM). After constructing a network, the genes with similar expression patterns were grouped into the same modules, and eigengenes were also calculated for these modules. Eigengene values were used to search the associations with saponins produced during fruit development. Then, *cis*-motif enrichment analysis of the TFs and saponin biosynthesis candidate genes in the chosen modules was conducted as described by Wang et al. (2021). First, the position frequency matrices (PFMs) of TFs were downloaded from plantTFDB (Tian et al., 2019); FIMO was then used to predict the *cis*-motif information in the promoter region (2000 bp upstream and 200 bp downstream of the transcriptional start site) of the saponin biosynthesis candidate genes under the condition *p* < 1e-3 (Grant et al., 2011). Finally, we constructed transcriptional regulatory networks by combining the availability of *cis*-acting element binding sites present in the promoters of the saponin biosynthesis candidate genes and the Pearson correlation coefficient (*r*>0.8) between these candidate genes and TFs in the same WGCNA module. In addition, the networks were visualized using the Cytoscape (v3.8.2) (Shannon et al., 2003).

### qRT-PCR analysis

Total RNA from pericarp at eight stages of growth was prepared according to the protocol described above. The first-strand cDNA was synthesized using a Goldenstar™ RT6 cDNA Synthesis Kit Ver 2 (Tsingke, Beijing, China) with reverse transcriptase. Eleven candidate genes and five TFs involved in saponin biosynthesis identified in the RNA-Seq data were selected for qRT-PCR analysis, and *SmACT (Samuk13G0061200)* was used as a reference to normalize gene expression. All primers used in qRT-PCR are listed in Supplemental Table S13. Reactions were performed on a LineGene 9600 Plus quantitative Real-Time PCR detection system (Bioer, Hangzhou, China) using a T5 Fast qPCR Mix (SYBR Green I) (Tsingke). Three biological replicates and three technical replicates were carried out for each stage. The relative expression level of each gene was determined by calculating the fold change in selected stages of pericarp samples relative to stage S1 using the 2^-ΔΔCt^ method.

### Yeast one-hybrid (Y1H) assays

Y1H assays were carried out based on the method described previously by Mou et al. (2018). Fragments of the target gene (*SmCYP71D-3*) promoter containing the core *cis*-elements were directly synthesized (Genecreate, Wuhan, China) and cloned into the pAbAi vector. The resulting pAbAi-*SmCYP71D-3* plasmids produced were then linearized and transformed into the Y1H Gold strain and selected with medium lacking Leu and supplemented with AbA (SD/-Leu/AbA medium). The open reading frames of TFs were inserted into the pGADT7 vector and transformed into Y1H Gold strains containing pAbAi-*SmCYP71D-3*. The empty pGADT7 vector and pGADT7-P53 were used as negative and positive controls, respectively. The transformants were further cultivated in medium lacking Leu (SD/-Leu medium). Positive yeast clones were grown on SD/-Leu liquid medium and diluted to different concentrations (OD_600_ = 10^-1^, 10^-2^, 10^-3^, 10^-4^), and aliquots of 10 μL yeast suspensions were plated on the SD/-Leu/AbA plates. All transformation and screening procedures were conducted twice.

### Electrophoretic mobility shift assay (EMSA)

The full-length coding sequences of *SmbHLH2*, *SmTCP4*, and *SmWRKY27* were amplified and inserted into the pGEX4T-1 vector to generate GST fusion proteins. *SmbHLH2-GST, SmTCP4-GST,* and *SmWRKY27-GST* were purified using a NucBuster™ Protein Extraction Kit (Cat. #71183; Merck KGaA, Darmstadt, Germany) for use in further EMSA experiments. The 26-nt probes of the promoter containing the putative *cis*-element binding sites of *SmCYP71D-3* were synthesized and labeled with biotin at the 5’ end (Genecreate). Mutant probes and unlabeled cold probes (20-, 50-, 100-fold concentration of unlabeled oligonucleotides) were utilized for competition experiments. The probes were incubated with the nuclear extract at room temperature for 30 min. The entire reaction mixture was run on a nondenaturing 0.5×TBE 6% polyacrylamide gel for 1 h at 60 V and 4°C, and then transferred onto Biodyne® B nylon membranes (Pall Corp., Ann Arbor, MI, USA) (Hellman and Fried, 2007). Signals were visualized with reagents included in the Efficient Chemiluminescence Kit (Vigorous, Beijing, China) and ChemiDoc XRS (Bio-Rad, Hercules, CA, USA).

### Statistical analyses

One-way analysis of variance (ANOVA) and Duncan’s multiple comparison test were used to analyze the differences in total saponin content at different stages using SPSS software (version 25.0; IBM Inc., Chicago, IL, USA). In all analyses, *p*<0.05 was taken to indicate statistical significance. Charts were plotted using Origin 2017 SR2 software (OriginLab Inc., Hampton, MA, USA). Correlation analysis, hierarchical clustering and principal component analysis (PCA) were performed using the corrplot and prcomp package in R.

## Accession numbers

Sequence data have been deposited in National Center for Biotechnology Information (NCBI) Sequence Read Archive (http://www.ncbi.nlm.nih.gov/sra) with the BioProject ID PRJNA784159.

## Supplemental data

**Supplemental Figure S1.** qRT-PCR verified the selected genes. Relative expression levels of qRT-PCR were calculated using *SmACT (Samuk13G0061200)* as a standard. Normalized gene levels in stage S1 were arbitrarily set to 1. Pearson correlation coefficients were calculated by comparing qRT-PCR and RNA-seq data for each gene across all samples.

**Supplemental Figure S2.** Number of DEGs between two tested groups (*Q*-value < 0.05 and |log_2_(Fold Change)| > 1). Here, FC refers to the log2 of the ratio of gene expression between two tested groups.

**Supplemental Figure S3.** GO enrichment analysis of DEGs between eight pericarp growth stages. Top 50 GO categories assigned to the DEGs. The genes were categorized based on gene ontology annotation and the proportion of each category is display in the categories of biological process (BP), cellular component (CC) and molecular function (MF).

**Supplemental Figure S4.** Temporal expression patterns of DEGs in the eight growth stages of the soapberry pericarps. All DEGs were classified into twelve clusters based on their expression patterns using the Mfuzz R package.

**Supplemental Figure S5.** Distribution of enriched KEGG pathways for various DEGs expression patterns in the pericarp growth. KEGG pathways involved in the biosynthesis of metabolites of cluster 2 (A), cluster 9 (B) and cluster 11 (C). Each circle represents a KEGG pathway.

**Supplemental Figure S6.** Phylogenetic tree of the soapberry *β-ASs* and *LUS.* The phylogenetic tree is constructed based on the deduced amino acid sequences for the soapberry *β-ASs* and *LUS* (bold letters) and other plant *β-ASs* and *LUS* involved in triterpenoid biosynthesis via the Neighbor-Joining method and established the reliability of each node through bootstrap methods, using ClustalW. Protein sequences are retrieved from NCBI GenBank using the following accession numbers: *Lotus japonicus LjAMY1* (AB181244.1), *LjAMY2* (AF478455.1) and *OSC3* (AB181245.1); *Glycyrrhiza glabra GgbAS1* (AB037203.1) and *GgLUS1* (AB116228.1); *Betula platyphylla* var. *japonica BPY* (AB055512.1) and *BPW* (AB055511.1); *Panax ginseng PNY1* (AB009030.1) and *PNY2* (AB014057.1); *Eleutherococcus senticosus EsBAS* (KX378998.1); *Barbarea vulgaris LUP2* (KP784688.1); *Arabidopsis thaliana AtBAS* (AB374428.1); *Vaccaria hispanica* VhBS (DQ915167.1); *Chenopodium quinoa CqbAS1* (KX343074.1).

**Supplemental Figure S7.** Phylogenetic tree of the soapberry CYP450s. The phylogenetic tree is constructed based on the deduced amino acid sequences for the soapberry CYP450s (bold letters) and other plant CYP450s involved in triterpenoid biosynthesis via the Neighbor-Joining method and established the reliability of each node through bootstrap methods, using ClustalW. Protein sequences are retrieved from NCBI GenBank using the following accession numbers: *L. japonicus LjCYP71D353* (KF460438.1); *Platycodon grandiflorus PgCYP716A141* (KU8878855.1); *Artemisia annua AaCYP716A14v2* (KF309251.1); *C. quinoa CqCYP716A78* (KX343075.1); *Aquilegia coerulea AcCYP716A110* (KU878864.1); *Kalopanax septemlobus KsCYP72A397* (KT150517.1); *Medicago truncatula MtCYP72A63* (AB558146.1) and *MtCYP72A68v2* (DQ335782.1); *Glycine max GmCYP72A69* (LC143440.1); *Glycyrrhiza uralensis GuCYP72A154* (AB558153.1).

**Supplemental Figure S8.** Phylogenetic tree of the soapberry UGTs. The phylogenetic tree is constructed based on the deduced amino acid sequences for the soapberry UGTs (bold letters) and other plant UGTs involved in triterpenoid biosynthesis via the Neighbor-Joining method and established the reliability of each node through bootstrap methods, using ClustalW. Protein sequences are retrieved from NCBI GenBank using the following accession numbers: *V. hispanica VhUGT74M1* (DQ915168.1); *Panax japonicus* var. *bipinnatifidus PjOAGT* (MH819287.1); *Panax zingiberensis PzOAGT1* (MH819284.1), *PzOAGT2* (MH819285.1) and *PzOAGT3* (MH819286.1); *Panax quinquefolius Pq3-O-UGT1* (KR028477.1) and *Pq3-O-UGT2* (KR106207.1); *Siraitia grosvenorii SgUGT74AC1* (HQ259620.1), *SgUGT720-269-1* and *SgUGT94-289-3; P. ginseng PgUGTPg45* (KM401918.1), *PgUGTPg100* (KP795113.1) and *PgUGTPg101* (KP795114.1).

**Supplemental Figure S9.** The correlation networks of genes in the blue (A) and greenyellow (B) modules, in which only edges with weight above a threshold of 0.35 and 0.45 are displayed, respectively.

**Supplemental Figure S10.** Transcriptional regulatory network of *SmCYP71D-3.*

**Supplemental Table S1.** Triterpenoid saponin metabolites tentatively detected in soapberry pericarps of different fruit growth stages.

**Supplemental Table S2.** Differentially accumulated saponins between different fruit growth stages.

**Supplemental Table S3.** The results of Mfuzz clustering for all of DASs in the eight fruit growth stages of the soapberry pericarps.

**Supplemental Table S4.** Summary of RNA sequencing and mapping using the soapberry genome as a reference.

**Supplemental Table S5.** List of saponin biosynthesis candidate genes.

**Supplemental Table S6.** Module sizes of WGCNA.

**Supplemental Table S7.** List of genes in blue module correlation network (with edge weight ≥ 0.35).

**Supplemental Table S8.** List of genes in greenyellow module correlation network (with edge weight ≥ 0.45).

**Supplemental Table S9.** Gene annotation in formations of blue module.

**Supplemental Table S10.** Network nodes of structural TF–gene network in the blue module.

**Supplemental Table S11.** Gene annotation in formations of greenyellow module.

**Supplemental Table S12.** Network edges of structural TF–gene network in the blue module.

**Supplemental Table S13.** Primers of sequences for qRT-PCR analysis.

**Supplemental Method S1.** Supplemental materials and methods.

## Acknowledgements

We sincerely thank Shuijing Luo for his tending of the plants. We are also grateful to Kunjing Wu and Kui Liu for his guidance in the experiment. We gratefully acknowledge the assistance of Xin Wang and Jing Zhong in this study.

